# Mango: Distributed Visualization for Genomic Analysis

**DOI:** 10.1101/360842

**Authors:** Alyssa Kramer Morrow, George Zhixuan He, Frank Austin Nothaft, Eric Tongching Tu, Justin Paschall, Nir Yosef, Anthony D. Joseph

## Abstract

The decreasing cost of DNA sequencing over the past decade has led to an explosion of available sequencing datasets, leaving us with terabytes to petabytes of data to explore and analyze. It is critical for analysts in research and clinical settings to be able to develop new data-driven hypotheses from these datasets through bias identification, analysis of data quality, and testing different algorithms and parameter settings. However, current interactive tools for sequence analysis are designed to run on single machines that do not scale to the size of modern genomic datasets, and rely on precomputed static views, rather than allowing direct interaction with the primary dataset. Mango is a genomic sequence visualization and analysis platform that removes these constraints regarding scalability and staticity by leveraging the power of multi-node compute clusters in the cloud to allow interactive analysis over terabytes of sequencing data. Mango provides both a genome browser graphical user interface and programmable notebook form factor to allow users of varying analytical experience to explore large sequencing datasets on both private clusters and in the cloud. These tools provide a flexible environment for interactive exploration of genomic datasets, while surpassing the computational limits of single-node genomic visualization tools.

## Introduction

The transition from Sanger to high-throughput DNA sequencing has led to a dramatic expansion of genomic sequencing datasets (Shendure & Ji 2008), exceeding the computational limits of single node genomics analytics tools (Schadt et al. 2010). While single whole genome sequencing samples have increased coverage depth resulting in more than 250 GB of reads, cohort studies have simultaneously targeted larger and more diverse populations, leading to studies that contain more than 300 TB of genomic variation data from a single cohort (Consortium et al. 2015, Consortium 2010, Mallick et al. 2016, Weinstein et al. 2011). Prior to the advent of cloud computing, working with these datasets required costly data transfers from centralized repositories like dbGaP (Zhang et al. 2008). However, cloud computing providers such as Google Cloud, Amazon Web Services, and Microsoft Azure have currently staged more than 600 TB of open access sequencing datasets into cloud stores, while initiatives like the Cancer Cloud Pilots have made sequencing projects like The Cancer Genome Atlas (TCGA) accessible via cloud storage (Lau et al. 2017, Weinstein et al. 2011). Simultaneously, big data distributed processing frameworks such as Apache Spark (Zaharia et al. 2010) and ADAM (Massie et al. 2013) have provided the necessary abstractions to read, process and manipulate terabyte sized genomic datasets in parallel in a cloud environment.

Together, the elasticity of the cloud and existing distributed systems tools provide the computational resources and APIs to interactively explore these massive datasets. However, tools for exploration and visualization of sequencing datasets are not yet optimized for cloud computing, and subsequently are limited to local analysis of only a few samples at a time. Additionally, existing tools tightly couple functionality with data layout, supporting a subset of queries and visualizations that have been pre-computed from the primary dataset. These constraints have not only restricted tools from taking advantage of cloud computing, but have also resulted in inherently static analyses, stifling the progression of exploratory data analysis in the field of genomics (Unwin 2008).

To address these challenges, we introduce the Mango browser and the Mango notebook: two tools which take advantage of elastic compute to allow exploration of personal and cloud staged datasets that surpass the computational resources of a single machine. Architecture for the browser and notebook are shown in Figure S1, and are built on Apache Spark (Zaharia et al. 2012) and ADAM (Nothaft et al. 2015), which enable fast parallel processing of terabyte-sized genomic datasets on a general purpose compute cluster. Both tools expose flexible query interfaces to empower users of various programming expertise to interactively explore datasets.

The Mango browser is a distributed genome browser, which provides a graphical user interface (GUI) for local analysis of genomic reads, variants, and features at user-defined loci, supporting functionality for quality analysis of sequencing samples, local correspondence between multiple samples, and further exploration of surrounding regions at interesting loci. The Mango browser supports functionality implemented in commonly used browsers such as IGV, IGB, JBrowse and Savant (Robinson et al. 2011, Freese et al. 2016, Fiume et al. 2010, Skinner et al. 2009) (Table S2), while scaling to dataset sizes surpassing the computational resources of single node tools (Figure S2). To achieve fast queries over genomic ranges, the Mango browser organizes data in distributed memory using Interval Resilient Distributed Datasets (Interval RDDs), a distributed implementation of interval arrays which support overlapping region queries in sub second latencies (see Supplementary 1.1.2). The Mango browser caches genomic regions using Lazy Materialization, which selectively caches genomic ranges that are sized based on the amount of memory available (see Supplementary 1.1.1).

The Mango notebook supports exploratory data analysis (EDA) in a Jupyter notebook environment (Kluyver et al. 2014), removing static constraints of genome browsers by allowing users to combine predefined and custom visualizations to summarize large data sets. Within a notebook, Mango provides the ability to visualize raw genomic reads, variants, and features around a single locus, as well as globally summarize data quality, mutation, and coverage distributions across multiple samples (Figure 2).

Both the Mango browser and notebook can leverage public datasets and elastic compute while achieving sub-second response times on cached datasets. While the Mango tools achieve comparable performance of traditional genome browsers like IGV when running on a single remote machine (see Supplementary 2), they surpass the functionality of existing tools by enabling analysis of multiple high coverage sequencing samples on a high performance computing cluster (Figures S2).

## Results

In the advent of big data analytics, data science workloads can be divided into two categories: discovery-driven decisions, and repeat, or cyclic, decision making (Provost & Fawcett 2013). We demonstrate the ability of the Mango browser and notebook to address data-driven and cyclic decisions by exploring data from the Illumina Platinum trios and the Simons Genome Diversity Project dataset (Mallick et al. 2016, Eberle et al. 2015), whose data sizes surpass the memory limitations of single node visualization tools on general purpose machines. We demonstrate the use of the Mango browser, along with ADAM (Massie et al. 2013), to drive de novo mutation discoveries from multiple high coverage sequencing samples and variants, while the Mango notebook supports cyclic decision making through global quality analysis of whole-genome sample coverage distributions, and corresponding local analysis on user-identified loci with high insertion rates. Both use cases demonstrate the novel functionality of interactive exploration on remotely staged datasets in a distributed cluster environment.

### The Mango browser enables discovery driven decisions through de novo variant exploration in the Illumina Platinum dataset

We use the Mango browser and ADAM to demonstrate a discovery-driven decision pipeline through the identification and exploration of missense de novo variants in the probands of the NA12877 and NA12878 trios from the Illumina Platinum dataset (Eberle et al. 2015), staged remotely on a general purpose compute cluster. Discovery and exploration of de novo mutations is a vital part of understanding causal effects of variants not selected for by evolution (Veltman & Brunner 2012), and requires both global and local analysis to identify genome-wide de novo mutations and subsequently verify specific loci with potentially causal mutations. The Mango browser and ADAM provide a complete end to end pipeline for querying and exploring such hypotheses, and reduce time to discovery through a single pipeline to answer their question of interest.

Figure 1 demonstrates visualization of de novo mutations in the Mango browser of the NA12877 and NA12878 trios. Using the Mango browser, users can visualize genome-wide de novo mutation sites on the Mango browser discovery page, and navigate to specific loci to visually verify sequence fidelity around sites with potential de novo mutations in the proband (Figure 1C). Figure S2 shows example query response times for alignment data for NA12878 from the Illumina Platinum dataset on 1093 virtual cores and 1TB memory. Interactive response times are reached when data is cached in memory and retrieved via Interval RDDs.

**Figure 1:**
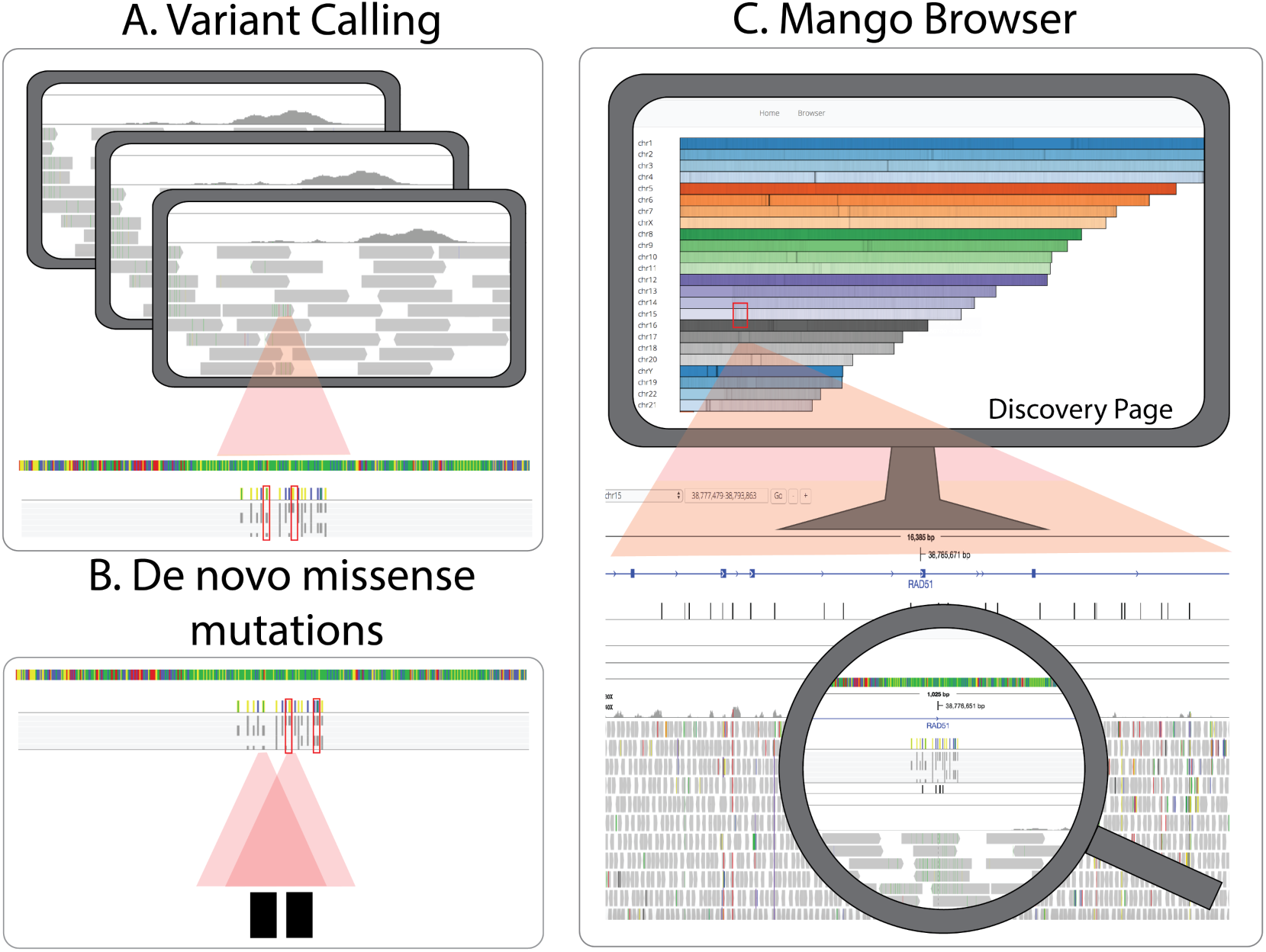
Six high coverage alignments totalling 692 GB were joint variant called using Avocado, a distributed variant calling tool built on the ADAM APIs (Nothaft et al. 2015, Nothaft 2017). Variant files were annotated using SnpEff. Resulting variants from Avocado were filtered for de novo variants in NA12877 and NA12878 and saved as ADAM feature files in Parquet format. Example query response times for the Mango browser are shown in Figure S2. **A.** Resulting variant calls and corresponding alignment data were loaded and visualized in the Mango browser. **B.** De novo missense mutations were observed through the variant track in the Mango browser. **C.** Global sites with de novo mutations could be visualized and selected on the discovery page and validated by visualizing raw alignment data.

### The Mango notebook supports exploratory data analysis on the Simons Genome Diversity Project dataset

We demonstrate the use of the Mango notebook to perform exploratory data analysis (EDA) through quality analysis of 12 high coverage samples from the Simons Genome Diversity Project dataset. EDA is becoming a crucial part of genome analysis, and is not driven by, but rather fosters hypotheses about the data (Behrens 1997). For EDA, users require a more flexible environment to iteratively query and analyze their data, allowing iterative discoveries to drive the next question of interest. For this use case, the Mango notebook supports a programming interface built on Jupyter notebook, allowing users to query, visualize and iterate through genomic analyses over remotely staged terabyte sized datasets. The Mango notebook supports visualizations for global and local analysis of genomic regions, as well as query support through Spark SQL (Table S1).

Figure 2 demonstrates the use of the Mango browser to identify outliers with low or high sequencing depth by generating coverage distributions over a population of 12 high coverage samples from the Simons Genome Diversity Project dataset (1.5 TB of alignment data). After summarizing coverage across samples, insertion distributions were generated on individual samples with high coverage distribution. Specific loci with high insertion distributions were then verified at single base pair resolution by viewing raw pileup data at specific genomic locations in the Mango browser. This global to local analysis of whole genome summaries to specific loci allows users to summarize multiple samples and validate fine-grained observations in a single environment, while being able to rapidly perform subsequent analyses driven by real-time observations.

**Figure 2:**
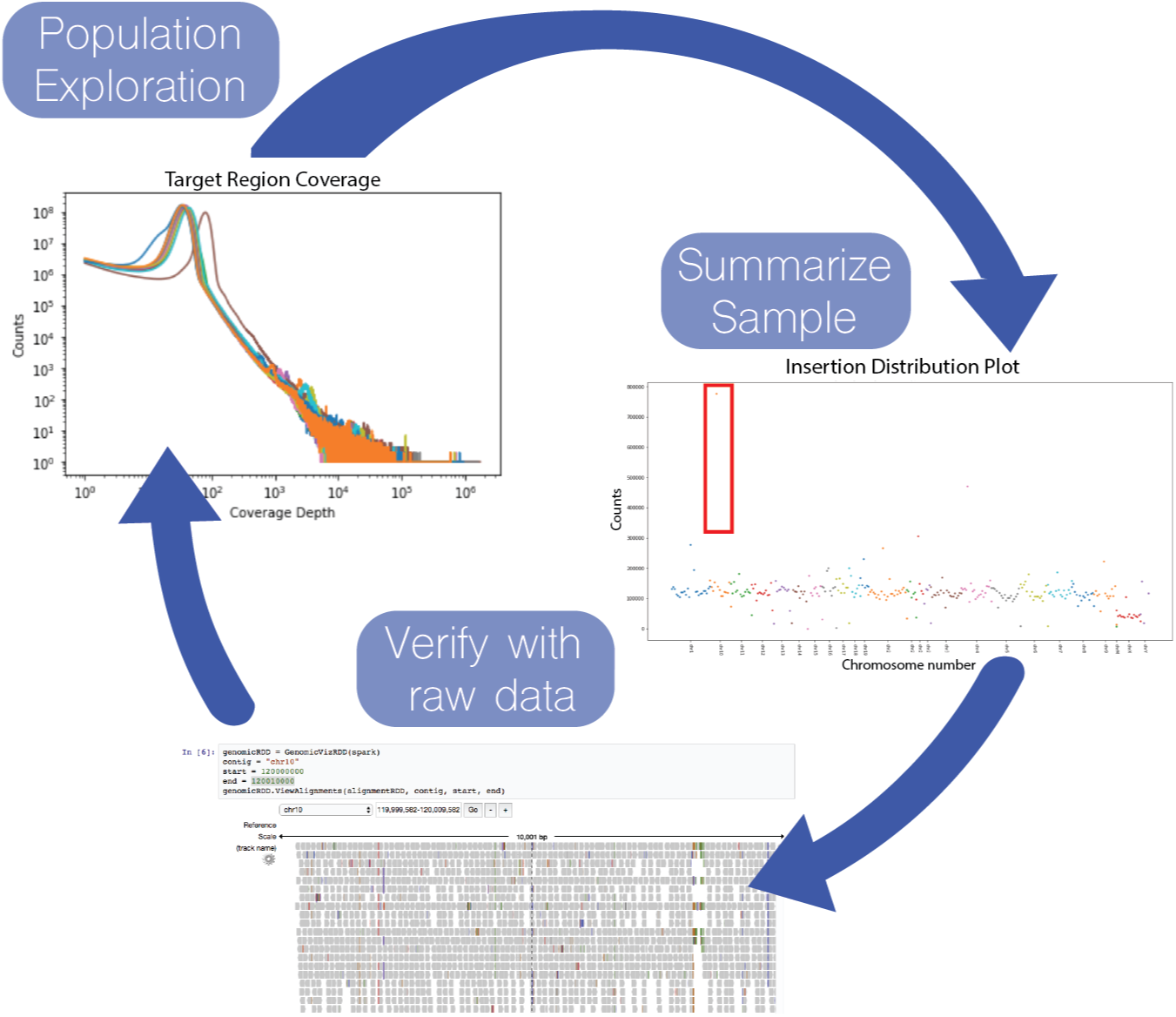
Cyclic analysis of 12 samples from the Simons Genome Diversity Project dataset in the Mango notebook. Coverage distributions were computed for 12 high coverage sequencing samples and plotted (top left). High coverage outlier in orange was selected for generation of insertion distribution plots (middle right). Sites of high insertion counts, outlined in red, were further explored by visualizing the raw alignment data for investigation of insertions on chromosome 10 (bottom). Table S1 lists additional functionality in the Mango notebook.

## Discussion

We have introduced the Mango browser and notebook, two visualization tools which enable EDA on large genomic datasets in a cloud computing environment. Both tools are built on Apache Spark, which supports efficient scaling to terabyte sized datasets, and ADAM, which provides support for genomic analyses on Apache Spark. While the Mango browser provides a GUI for exploring genomic datasets that scales past single node genome browsers, the Mango notebook provides a programmable environment that allows a tight loop between customized queries and common visualizations that accelerates the exploration of genomic datasets. Both tools remove pre-processing and scaling constraints of current genome visualization tools, allowing users to browse heterogeneous datasets from multiple platforms as they are staged by cloud providers, and scale past the computational limits of a single machine.

## Methods

### Contact for Reagent and Resource Sharing

Further information and requests for resources should be directed to and will be fulfilled by the Lead Contact, Alyssa K. Morrow.

## Method Details

Unless otherwise noted, all performance evaluations were run on our internal cluster, which is comprised of 55 compute nodes, each with two eight-core Intel Xeon E5-2670 CPUs with 2x hyperthreading, 256 GB RAM and 4TB of HDD (four 1TB 7200 RPM HDDs connected via 6Gpbs SATA). The nodes are connected by a Gigabit ethernet network. The 4TB of HDD per node is used as backing storage for an Apache Hadoop Distributed File System (HDFS), (*The Hadoop Distributed File System* 2010) deployment running at 2x replication with a total configured capacity of 190TB. Additionally, a 30TB Pure Storage FlashBlade is mounted across all nodes in the cluster. The FlashBlade is connected to all nodes by a separate 10GbE network. Jobs are submitted to the cluster using the Apache Hadoop YARN scheduler (*Apache Hadoop YARN: yet another resource negotiator* 2012). A virtual core represents a single hyperthreaded instance of a physical core, thus there are twice as many virtual cores as physical cores in the cluster.

### Processing the Illumina Platinum dataset

#### File Conversion and Variant Calling

Six whole genome files totalling 691 GB from the Illumina Platinum dataset were staged in HDFS and converted from BAM to ADAM format. Total conversion time was 72m (minutes), averaging 12m per file. Final total size for converted alignment files was 822 GB. These six files were joint variant called using Avocado, a distributed variant calling tool (Nothaft et al. 2015, Nothaft 2017). Variants were annotated for missense mutations using SnpEff (Cingolani et al. 2012).

#### Filtering variants in the proband

The resulting variant calls we queried for de novo variants in the proband for NA12877 and NA12878 trios using Spark and ADAM. These queries ran in 7m27s (seconds) and 4m54s for NA12877 and NA12878, respectively. The resulting regions with de novo variants were saved as a Feature RDD and loaded into Mango. In conjunction with Mango’s discovery mode, all de novo regions were preloaded into the browser. NA12877 had 90705 regions suggesting de novo variants. NA12878 had 85058 regions suggesting de novo variants. De novo regions were then filtered for missense variants using ADAM’s join functionality. We found 16 de novo missense mutations for NA12877 and 11 de novo missense mutations in NA17878.

#### Viewing variants in the Mango Browser

Six alignment files, de novo regions and variants were loaded into the Mango browser. Users can load either indexed BAM files or converted Parquet files into the browser, although loading an indexed BAM file incurs extra start up latency (Figure S2). Figure S2 shows the latencies from using these different file access modalities. Other configurations for the Mango browser include discovery mode and partitioned Parquet files. Partitioned Parquet files speed up region queries by sharding large files by chromosome and genomic start position. This allows Parquet to only scan a subset of blocks when initially loading genomic ranged data, decreasing initial latency by 10x compared to non-binned Parquet files (Figure S2). Discovery mode preloads variants and features, allowing users to visualize and quickly explore such datasets when traversing new regions in the browser. Loading time for feature and variant data with and without discovery mode follow similar query patterns as alignment files in Figure S2. Although discovery mode incurs an 7x initial latency during setup, latency for subsequent requests are reduced by 80-90x, achieving interactive latencies of less than 500ms for both variants and features.

### Simon Genomes Diversity Dataset analysis

#### Alignment of samples

We used Cannoli (bdgenomics 2018) and BWA MEM (Li 2013) to align reads from the Simons Genome Diversity Dataset (SGD) to the GRCh38 reference. Cannoli uses ADAM’s pipe API to run BWA MEM in parallel across a cluster using Apache Spark. ADAM’s pipe API executes a bioinformatics tool as a subprocess on the Apache Spark executor and translates the data between Apache Spark’s in-memory representation and the common file formats used by bioinformatics tools. This allows alignment using BWA MEM to run in 11m on a 1,024 core cluster for a 50x coverage WGS sample. After alignment, ADAM was used to realign INDELs with default parameters. The Toil (Vivian et al. 2017) scripts used to run this pipeline are available from the bdgenomics.workflow package at github.com/bigdatagenomics/workflows.

#### Coverage Conversion

12 SGD samples were converted to coverage files using ADAM’s toCoverage function. Average conversion time for the 12 samples was 8m49s.

#### Coverage Distribution and Indel Plots

Coverage distributions were generated and visualized in the Mango notebook using 1721 virtual cores and 968 GB memory. Total time for coverage plot generation shown in Figure 2 was 10m14s. Insertion distribution for a single file was 3m29s. Deletion distribution was instantaneous, as information was aggregated and cached from the previous insertion distribution plot. Alignments from regions in the distribution plots were filtered and viewed with a pileup, taking 18 seconds and 20 seconds to view regions of size 1,000 and 10,000 base pairs. respectively.

## Data and Software Availability

### Dataset Availability

The datasets analyzed in this paper include the Illumina Platinum dataset (Eberle et al. 2015) and the Simons Genome Diversity dataset (Mallick et al. 2016). The Illumina Platinum dataset is available through BaseSpace Sequence Hub and the European Nucleotide Archive, with ENA accession PRJEB3381. The Simons Genome Diversity dataset is available through the EBI European Nucleotide Archive under accession numbers PRJEB9586 and ERP010710.

### Code Availability

Mango is Open Source software and is released under a permissive Apache 2 license. Mango follows a community-oriented development model and will expand features and available visualizations based on community requests. Source code and example files for Mango is available from https://github.com/bigdatagenomics/mango. Documentation is available at http://bdg-mango.readthedocs.io/en/latest/, which contains tutorials and access to distribution code.

## Additional Resources

The following configurations and their documentation exist for running the Mango notebook and browser:

### Building from source with examples

http://bdg-mango.readthedocs.io/en/latest/installation/source.html

### Running Locally from Distribution Code

http://bdg-mango.readthedocs.io/en/latest/installation/distribution.html

### Running Locally through Docker

http://bdg-mango.readthedocs.io/en/latest/docker/docker-examples.html

### Running on Amazon Elastic Map Reduce

Instructions for running the Mango browser and notebook on Amazon Elastic Map Reduce (EMR, https://aws.amazon.com/emr/) can be found at http://bdg-mango.readthedocs.io/en/latest/cloud/emr.html. This tutorial includes hardware requirements and cluster configuration requirements. Example files for EMR can be found in the Mango Docker container at /opt/cgl-docker-lib/mango/example-files/notebooks/aws-1000genomes.ipynb. Initial configuration time, including AWS configuration, EMR cluster creation and bootstrapping time is approximately 1 hour.

### Running on Google Cloud Engine

Instructions for running Mango on Google Cloud Engine (https://cloud.google.com/compute/) can be found at http://bdg-mango.readthedocs.io/en/latest/cloud/google-cloud.html. Example files for GCE can be found in the Mango Docker container at /opt/cgl-docker-lib/mango/example-files/notebooks/gce-1000genomes.ipynb.

